# FAK Differentially Mechanoregulates Cell Migration During Wound Closure

**DOI:** 10.1101/2025.04.01.646098

**Authors:** Jennifer Patten, Nourhan Albeltagy, Jacob D. Bonadio, Armando Ortez, Karin Wang

**Author notes:** Corresponding Author: Karin Wang, 1947 North 12^th^ Street, Philadelphia, PA 19085.

## Abstract

Cell migration is an essential step in wound healing. Mechanical input from the local microenvironment controls much of cell velocity and directionality during migration, which is translated into biochemical cues by focal adhesion kinase (FAK) inside the cell. FAK induces both regeneration and fibrosis. The mechanisms by which FAK decide wound fate (regenerative or fibrotic repair) in soft, normal wounds or stiff, fibrotic wounds remains unclear. Here we show that FAK differentially mechanoregulates wound behavior on soft substrates mimicking normal wounds and stiff substrates mimicking fibrotic wounds by converting mechanical substrate stimuli into variable cell velocity, directionality, and angle during wound healing. Cells on soft substrates migrate slower and less persistently; cells on stiff substrates migrate faster and more persistently with the same angle as the cells on normal wound substrates. Inhibition of FAK results in substantially slower, less persistent, and less correctly angled cell migration, which leads to slowed wound closure. Moreover, FAK inhibition impairs fibroblast ability to respond to substrate stiffness when migrating. Here we show FAK is an essential mechanoregulator of wound migration in fibroblast wound closure and is responsible for controlling cell migration dynamics in response to substrate stiffnesses mimicking normal or fibrotic wounds.

TOC Graphic:
Cell migration dynamics regulated by substrate stiffness via focal adhesion kinase (FAK). Cells on softer substrates mimicking normal wounds display slower, more random migration that is impaired by FAK inhibition. Cells migrate faster and more persistently on stiffer substrates mimicking fibrotic wounds that is dysregulated by FAK inhibition.

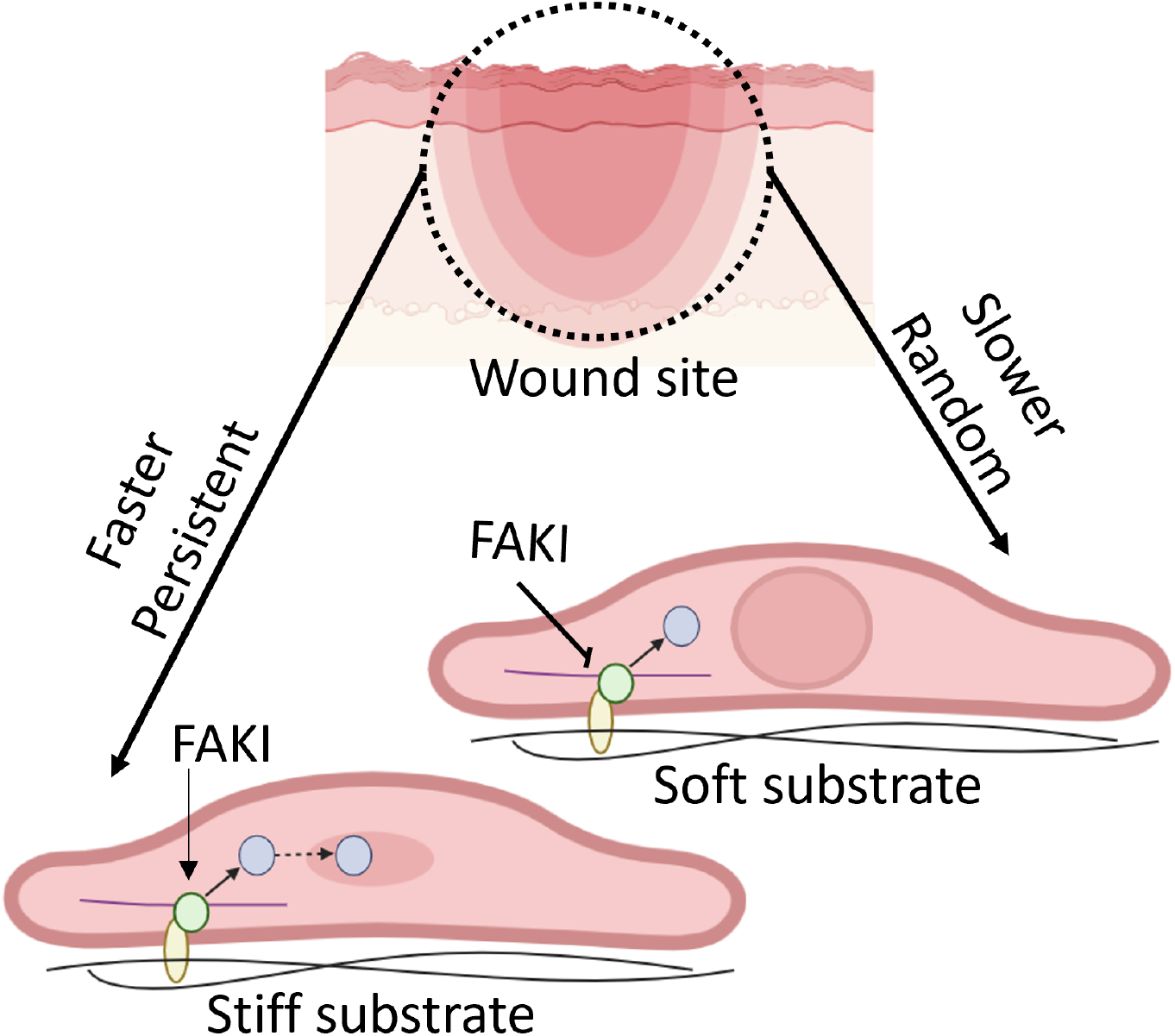

## Introduction

Cell migration is widely regarded as the rate limiting step in wound healing.^1–4^ Cells migrate in response to various stimuli, including chemotaxis (chemical attractants), durotaxis (mechanical signals), and gene regulation (biological drivers).^5–10^ Of these, mechanical forces regulate wound outcomes on multiple levels. Force mitigation is recognized to reduce scar formation by decreasing the fibrotic deposition of excess extracellular matrix. Force mitigation can even induce regeneration by restoring original tissue structure-function. Wounds bisecting Langer lines, a 3D map of mechanical load along the human dermis from gravity, location, and underlying extracellular matrix (ECM) structure, develop increased scar response,^11–13^ whereas areas of minimal to non-existent contraction (fetal cutaneous wounds,^14^ wound areas with no contractile forces^15^) undergo actual regeneration. Therefore, force mitigation is an important aspect of wound healing that determines which pathways are activated: fibrotic repair or regenerative repair.

Mechanistically, cells migrate into the wound by connecting to their local microenvironment via actin dynamic structures, lamellipodia and filopodia, and contracting to pull the cell body forward.^16^ On the interior of the cell, the focal adhesion protein complexes assemble to connect transmembrane receptor proteins to the actin cytoskeleton, which drives cell motility.^17^ Focal adhesion recruitment dynamics remain a consistent focus of study as the focal adhesion complex is subject to many regulators. The strength, size, and maturity of the focal adhesion complex is strongly affected by receptor-specific activation (ECM or cell binding) and mechanical input from local microenvironmental forces, leading to complex and nuanced activation profiles.^7,18,19^

Focal adhesion kinase (FAK) is a non-receptor protein tyrosine that is part of the focal adhesion complex.^20^ FAK is especially mechanosensitive and controls several mechanotransduction pathways.^17,21–24^ It senses mechanical signals from the local microenvironmental and translates them into biochemical cues driving cell migration and fibrosis.^25^ Substrate properties affecting FAK signaling include whether the model is 2D or 3D, soft or stiff,^26^ viscoelastic or elastic, ^26–28^ and can lead to various and non-linear FAK-mediated cell responses.^29–31^ Moreover, FAK has a complex and nuanced signaling profile that is dependent on many factors including cell density, cell area, presence/absence of downstream signaling effectors, and focal adhesion size (strength of connection to the local microenvironment).^19^ FAK can be directly correlated with the fibrotic response.^9,25,32^ During fibrosis, the extracellular matrix is stiffer and more aligned, providing a vastly different microenvironment for cells to interact with than that in wounded tissue.^33–35^ In 2D wound assays, FAK is necessary for wound contraction (closure).^9^ Moreover, inhibition of FAK directly promotes tissue regeneration.^36^ How FAK is active in both fibrotic repair and regenerative repair is an open question.

Migration is an essential step of wound healing; moreover, the rate of wound closure serves as a prognostic indicator of beneficial wound outcomes, including regeneration. The microenvironment plays an essential role in deciding wound fate by controlling various cell behaviors, including migration.^37,38^ FAK is an important element of the focal adhesion complex which connects the exterior cell environment to the internal cell signaling mechanisms. Why FAK is active in both fibrosis and regeneration is unclear. Our research question focused on whether we could induce a regenerative wound closure outcome by inhibiting key biochemical factors responding to fibrotic mechanical stimuli. Therefore, we investigated therapeutically targeting FAK’s mechanoresponses to inhibit fibrotic mechanical cues would restore normal wound closure.

## Materials and Methods

### Substrate Manufacturing and Cell Culture

The 2D wound assay polydimethylsiloxane (PDMS) substrates were fabricated as previously reported, which mimics the viscoelasticity of human skin at 18kPa and 146kPa by varying the base:crosslinker ratio appropriately (49:1, 40:1, respectively).^33^ Adult human dermal fibroblasts (HDFa, PCS-201-012 ATCC) of passage 8 or lower were seeded on 30µg/mL plasma fibronectin-coated PDMS substrates at 65,000 cells/100mm^2^ 24h prior to assay start. Cells were cultured in DMEM with 10% fetal bovine serum and 1% penicillin/streptomycin around a PDMS mask which, when removed, simulated wounding and began the wound assay.

### Inhibitor Conditions

FAK inhibitor FAKI (HY-10459, PF-562271, FAKI vs6062, Med Chem Express) was exogenously added into the cell culture medium at the start of the wound assay. The FAKI concentration was 5µM/mL.^36^ As DMSO was the solvent used for the inhibitor, a DMSO control of 5µM/mL was used to match the concentration of inhibitor used.

### Time Lapse Microscopy and Cell Analysis

Using a Keyence BZ-X8000 fluorescent light microscope, time lapse images of phase and Cy5 LED channels were taken with a 10x objective every 10 minutes for 48h post wounding. The individual .tiff images were then compiled to create time lapse image stacks. Inclusion criteria included no visible defects to the PDMS at the wound site and wound edges within the field of view. Time lapse image stacks were excluded if a significant amount of cells had migrated into the wound site under the PDMS mask prior to mask removal. The wound area was measured every 12h from the phase videos to quantify wound area over time (0,12,24,36,48h time points). The wound area was freehand traced in ImageJ Fiji around the cells migrating into the wound, and the area within the enclosed boundary was measured at each time point. Wound closure over time for all time slices were then divided by the original wound area (100%) measured at time 0h to return percentage of wound area closed. The equation for wound area is: (T_I_ /T_0_) x 100, where T_I_ is the wound area at a given time point, and T_0_ is the wound area at the wound start.

The Cy5 LED channel time lapse videos of wound fluorescent nuclei, labeled with SPY595 DNA tracker, were run through ImageJ Fiji TrackMate analysis to generate spots, identify fluorescent nuclei per time slice, and tracks, which combine the spots across all slices into a coherent single cell migration track.^39^ TrackMate spot sizes were limited to spots under 20µm. TrackMate duration track filters (excluding tracks with less than 4 spots) were applied to limit noise from artefacts and single spot recognition; the spot gap closing algorithm was limited to 4 time slices with 10 minutes elapsing between these slices, and less than 30µm distance traveled during that period to generate accurate tracks of cellular migration without overlap or confusion of tracks. The TrackMate tracks were further analyzed through Chemotaxis and Migration tool (Ibidi) to plot the migration data, including velocity, directionality, and cell angle. Directionality was taken as a measure of cell persistence, or consistent migration in the same direction. Therefore, a straight line is considered a 1, whereas a cell track with many switchbacks continually turning back on itself, or randomly migrating, is taken as a 0.

### Statistical Analysis

Sample size per condition was run with 3-5 separate wounds per condition. One-way ANOVA with Tukey’s post-hoc analysis or two-way ANOVA with Šídák’s multiple comparisons post-hoc analysis in GraphPad Prism 9 were used to evaluate each test condition as appropriate. Within each time point and test condition, cell responses on soft (normal) and stiff (fibrotic) substrates were compared. Across time points and test conditions, only similar stiffnesses were compared. Wound closure analysis was plotted with a best fit linear regression line. The slope and standard error of the best fit lines plotting the wound closure rate per condition were compared. All data presented as median ± standard deviation unless otherwise noted.

## Results and Discussion

### Substrate stiffness did not alter wound closure rate but did alter individual cell dynamics

We examined cell dynamics (Fig 1A,B,C) on soft or stiff substrates mimicking normal (18 kPa) and fibrotic (146 kPa) wounds, respectively. We observed that wound closure rate on softer, normal wound substrates was not significantly different compared to wound closure rate on stiffer, fibrotic wound substrates (Fig 1D). However, cells on softer substrates (normal wound) migrated significantly slower than cells on stiffer substrates (fibrotic wound) (Fig 1E). Cells on softer, normal wound substrates also migrated more randomly than cells on the stiffer, fibrotic substrates (Fig 1F). The axis of wound closure falls along the x-axis of 0-180° (Fig 1C,G). We observed that cells did not significantly change their angle of migration towards wound closure based on their substrate stiffness.

**Figure 1.**
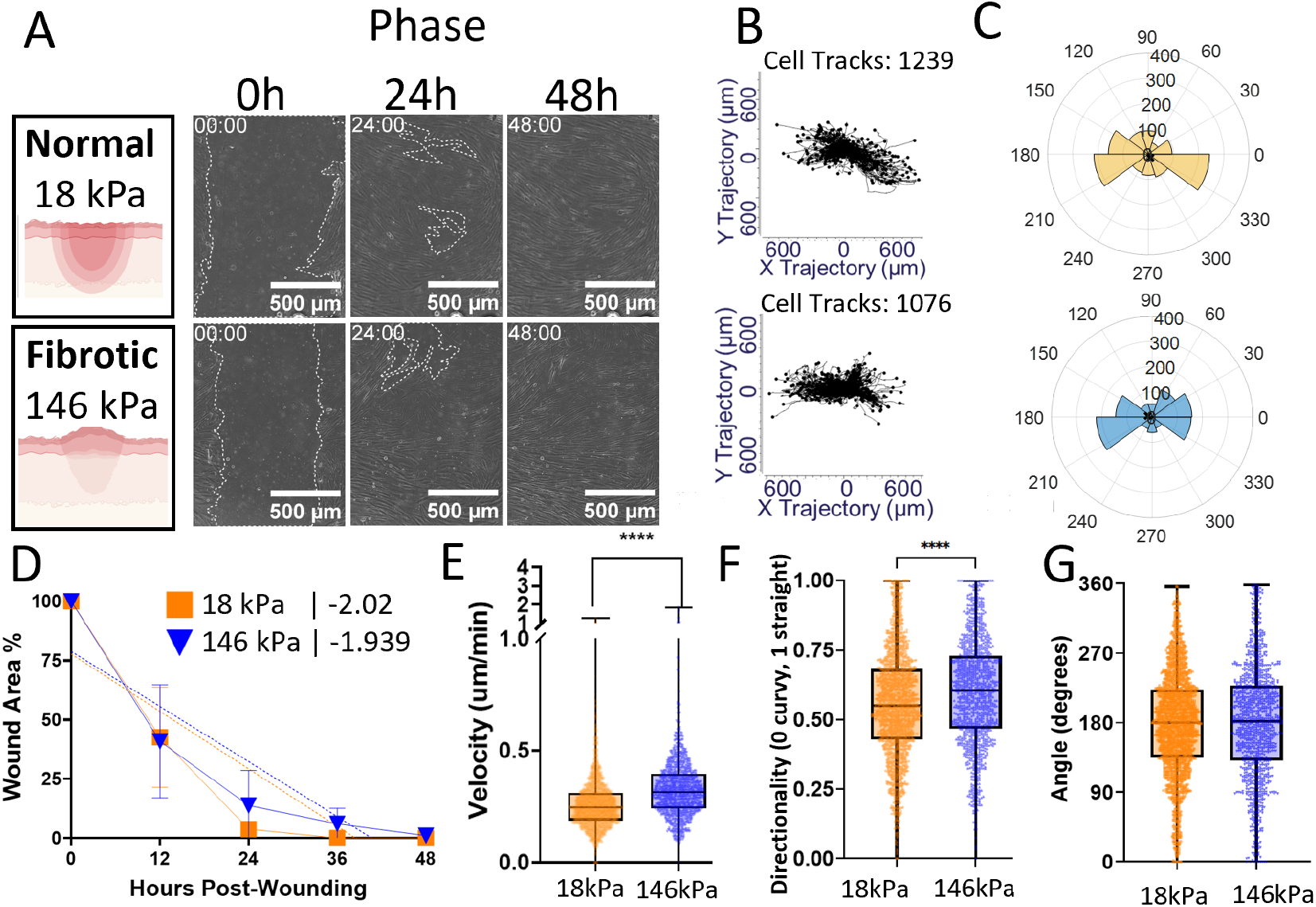
Cells on softer substrates mimicking normal wounds migrate slower and less persistently than cells on stiffer substrates mimicking fibrotic wounds. **A**. Representative 10x images of the wound on normal and fibrotic substrates, scale bar 500μm in phase microscopy, used to quantify wound area. In parallel, the time lapse videos were analyzed in TrackMate (ImageJ Fiji plugin) and Ibidi Chemotaxis and Migration Tool to extract cell migration dynamics, plotted as (**B**) representative cell tracks. **C**. Cell angle measured between start and end position. The angle of cell migration shown in degrees along four Cartesian quadrants, numbers shown around the plot perimeter. Radial numbers indicate number of cells per direction. **D**. No significant difference in rate of wound closure between substrate stiffnesses was observed. Linear regression analysis is plotted in dotted lines, and the slope is included in the legend. **E**. Cells on softer substrates migrated significantly slower than cells on stiffer substrates. ****P<0.0001. **F**. Cells migrated more randomly, or with less directionality and persistence, on softer substrates than cells on fibrotic substrates. ****P<0.0001. **G**. Cell migration angle towards the axis of wound closure does not change significantly based on substrate stiffness. N≥3 replicates. All significances evaluated with 2-way ANOVAs, with Šídák’s multiple comparisons post-hoc analysis.

In our model, cells behave according to the conventional paradigm that cell speed is greater on stiffer substrates.^40–42^ This relationship is mutable; depending on material surface properties and biochemical cues, cell speed on stiffer substrates can vary; these factors can also alter other migration parameters, including persistence.^43,44^ In our wound assay, we observed that persistence was greater (less random) on increased substrate stiffness. This is corroborated by existing literature that substrate stiffness coupled with an homogenous Fn coating leads to higher persistence.^43^ Moreover, our results also supports that wound closure rate is not dependent on substrate stiffness,^45^ though substrate stiffness does alter certain individual cell migration dynamics such as cell velocity and persistence.

Having established the dynamic cell responses to substrate stiffness in our model, we next examined how cells used FAK-mediated signaling to control the varying migration dynamics observed in response to mechanical stimuli from the substrate stiffness designed to mimic normal or fibrotic wounds.

### Inhibition of FAK slows wound closure rate inversely dependent on substrate stiffness

With the introduction of the FAK inhibitor, FAKI, we used DMSO as a control for comparison since our inhibitor was reconstituted in DMSO. The rate of wound closure (Fig 2A) from DMSO-treated fibroblasts was again not affected by substrate stiffness (Fig 2B). However, when FAK was inhibited on both soft and stiff substrates, fibroblasts slowed wound closure. In fact, when inhibiting FAK, wound closure on the stiffer substrate was the slowest, even slower than wound closure on the soft substrate. This suggests that FAK inhibition impairs cells’ ability to coordinate their internal contractile machinery to respond to the mechanical input from the microenvironment. Therefore, our data shows that FAK inhibition alter the cell’s ability to close the wound.

**Figure 2.**
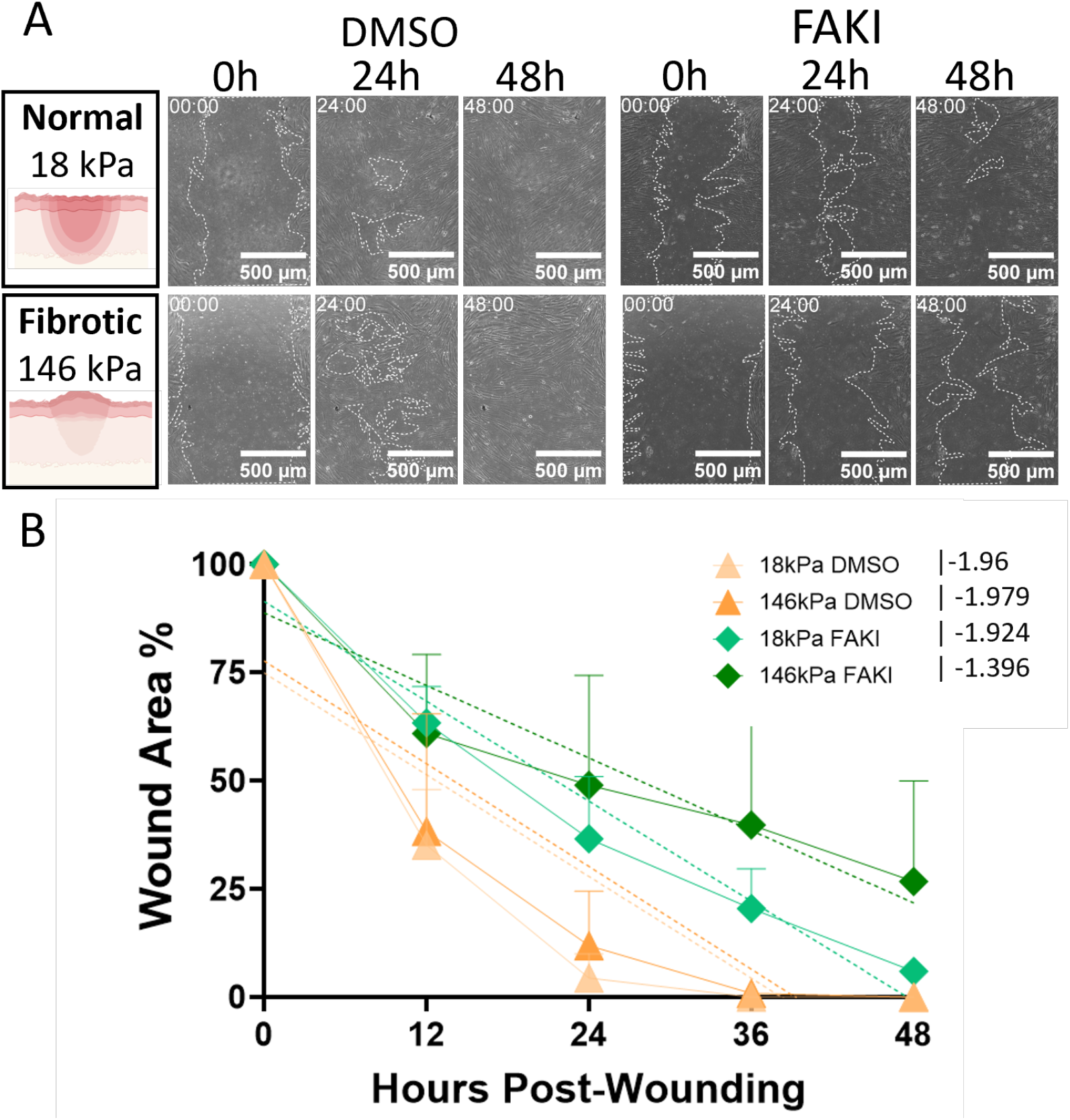
Inhibition of FAK slows wound closure rate inversely to substrate stiffness. **A**. Representative phase images at 10× objective. The wound area is marked by dashed lines, with the scale bar at 500μm. **B**. Wound closure is plotted as a percentage of wound area remaining. FAK inhibition slows gap closure time, indicating this protein is essential during wound closure. N≥3 replicates. Mean plotted as data points and standard deviation plotted in bars. Linear regression analysis is plotted in dotted lines, and the slope is included in the legend.

The DMSO-treated fibroblasts did not close the wound differently than the media control, regardless of substrate stiffness (Fig S1, S2A). The role of FAK in mediating wound closure remains unclear, with conflicting results appearing in the literature. Inhibition of FAK within *in vitro* models promote wound closure by attenuating the fibrotic response.^46^ Conversely, FAK inhibition also slows fibroblast migration (supported by our results).^21^ Within *in vivo* models, FAK inhibition leading to more wound closure, likely through less cell contractility.^47^ However, FAK promotion increases migration through α-catenin-mediated activation of FAK,^21^ pro-migratory FAK/ERK 1/2/YAP pathway,^48^ or by FAK inhibition by decreasing the fibrotic regulator FRNK.^32^ While this might suggest that activating the FAK-mediated pathway controls the nature of the cell response, the reality is not so clear. FAK can activate multiple types of responses through the same pathway. For example, FAK is known to regulate pro-migratory pathways, including PI3K, and ERK 1/2.^21,49^ Additionally, the integrin-based FAK-Src-PI3K-YAP pathway inhibits the Hippo pathway, resulting in decreased contact inhibition.^50^ In wound healing, α-catenin increases FAK/YAP activation to drive fibroblast migration and proliferation.^21^ However, FAK also regulates pro-fibrotic mechanisms through these pathways as well: ERK,^36^ and YAP,^51,52^ as well as others like the Akt pathway.^53^

Why various pathways – or even the same pathway - are activated within wound repair leading to either fibrosis or regeneration is unclear; how does mechanical stiffness interplay with FAK to regulate migration dynamics? Therefore, we quantified more nuanced individual cell migration dynamics to probe how FAK drives wound closure mechanisms in the context of soft and stiff substrates mimicking normal and fibrotic wounds, respectively.

### FAK inhibition slows cell migration

We next quantified fibroblast velocity to investigate migration dynamics (Fig 3A) behind the slower wound closure observed when inhibiting FAK signaling. Interestingly, FAK-inhibited fibroblasts demonstrated no significant differences in velocity between substrate stiffnesses (Fig 3B). Additionally, when FAK was inhibited, cell velocity on both substrate stiffnesses was significantly slower than their DMSO control counterparts. The addition of DMSO significantly slowed fibroblast velocity on softer substrates (Fig S1D).

**Figure 3.**
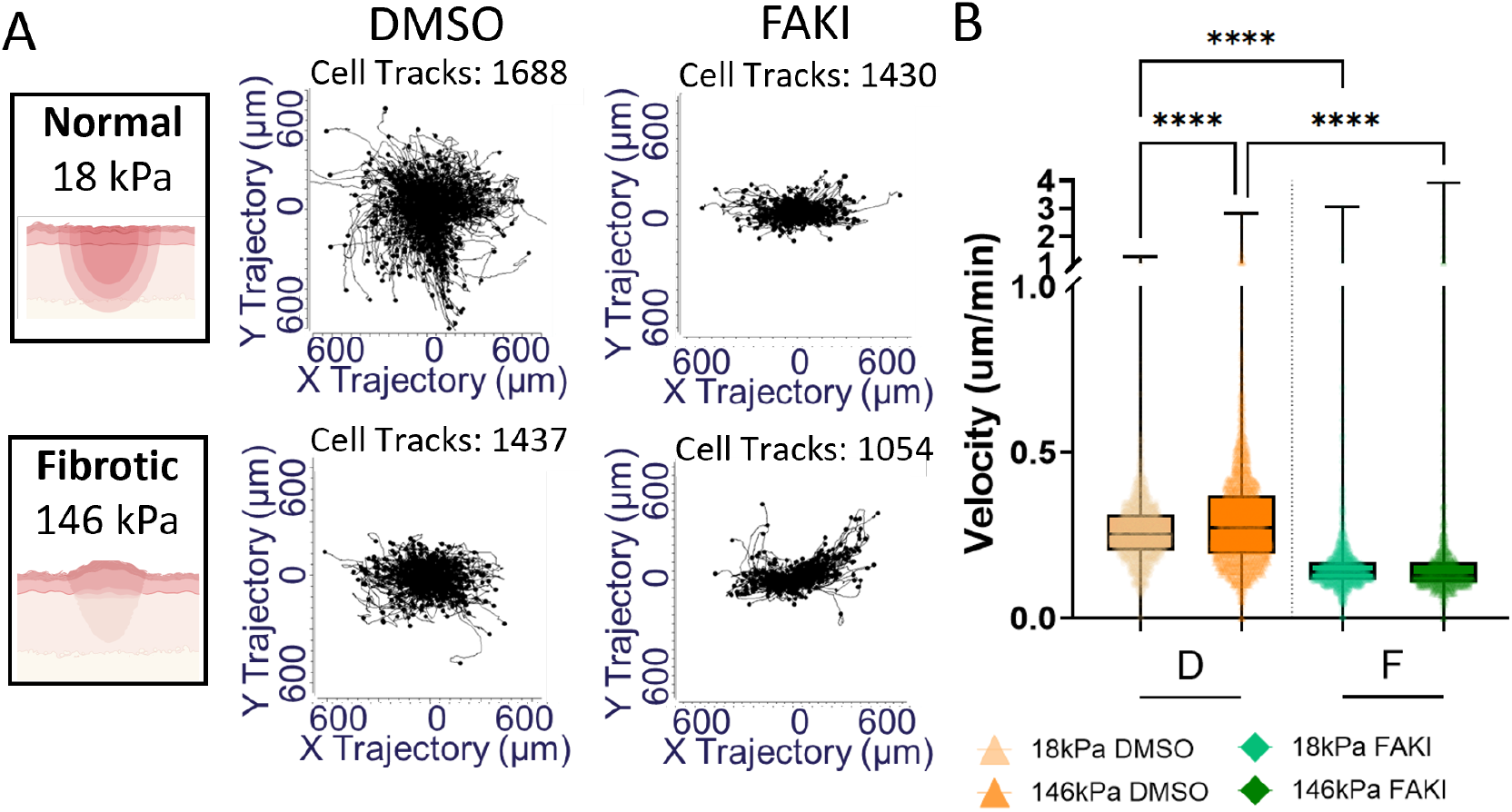
FAK inhibition slows cell velocity. **A**. Representative cell tracks per condition as plotted in Chemotaxis and Migration tool with data obtained from Track maté analysis of cell dynamics. **B**. Cell velocity obtained from Chemotaxis and Migration Tool after TrackMate analysis. ****P<0.0001. N≥3 replicates. All significances evaluated with 2-way ANOVAs, with Sidak’s multiple comparisons post-hoc analysis.

Our results indicate that in our fibroblast model, a low dose of DMSO (<0.005%) lowers fibroblast velocity compared to an untreated control. Previous work has indicated DMSO treatment in wound environments non-significantly reduces human dermal fibroblast migration compared to an untreated control.^54^

More importantly, FAK inhibition significantly slowed migration velocity regardless of substrate stiffness. This is consistent with literature, which indicates FAK, mediated by the TYR-397 site, is largely responsible for bulk migration response to mechanical input.^20^ FAK inhibition completely dysregulated the fibroblasts’ ability to change speed in response to varying substrate stiffnesses. However, the ability of a cell to continue migrating in a certain direction might also be impacted by mechanical stiffness cues. Therefore, we next studied cell persistence to shed light on the mechanisms of FAK-mediated migration dynamics.

### FAK mediates cell persistence

For migration to occur, a cell must be traveling along a trajectory (Fig 4A). Persistence is defined as the ability of the cells to continue migrating in one direction (Fig S2C). A cell that travels in a straight line is highly persistent while a cell that changes directions many times is not persistent and migrating randomly. The DMSO-treated fibroblasts migrated more randomly (less persistent) on the stiffer substrates (Fig 4B). Comparatively, DMSO-treated cells on the softer substrates mimicking normal wounds migrated with more persistence. This indicates that despite the faster migration velocity observed in Fig 1F, DMSO-treated cells unable to persistently migrate in the same direction on the stiffer substrates mimicking fibrotic tissue (Fig 3B). Compared to the original media conditions, DMSO treatment significantly downregulates cell persistence on stiffer substrates (Fig S2E). Additionally, treatment with DMSO promoted cell persistence on softer wound substrates compared to the media control (Fig S2E). The effect of DMSO on fibroblast persistence is not well studied. In this model, our data indicates DMSO deregulates fibroblasts’ ability to respond to their environment by promoting persistence on softer substrates and inhibiting this persistence on stiffer substrates.

**Figure 4.**
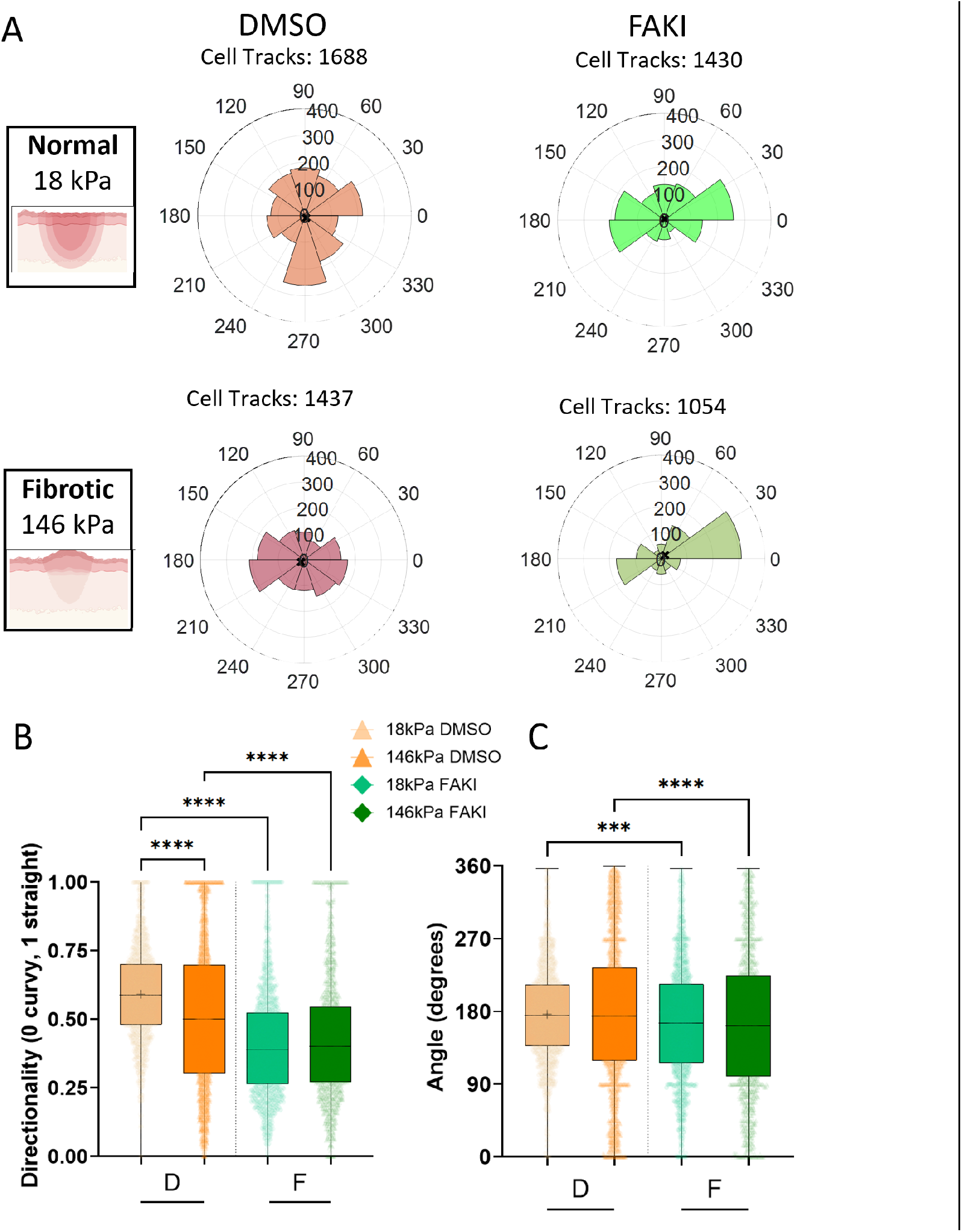
FAK inhibition dysregulates cell directionality and angles away from axis of closure. **A.** Representative rosette plots of cell angle during wound closure. In-house Matlab code used to plot Chemotaxis and Migration Tool-calculated angles. **B**. Cell directionality, ****P<0.0001. **C**. Cell angle measured between start and end position. The angle of cellular migration shown in degrees along four Cartesian quadrants, numbers shown around the plot perimeter. Radial numbers indicate number of cells per direction, ***P<0.0005, ****P<0.0001. N≤3327 cells from N≤3 wells plotted in median and standard deviation, two-way ANOVA, Šídák’s multiple comparisons post-hoc analysis.

FAK-inhibited fibroblasts on softer substrates showed significantly lower persistence compared to the DMSO control (Fig 4B). Likewise, on the stiffer substrates, cells treated with FAKI were significantly less persistent than their DMSO control counterparts. Moreover, unlike the DMSO controls, FAK inhibited fibroblasts showed no significant changes in cell persistence when migrating on soft (normal) and stiff (fibrotic) substrates. This indicates that the mechanism by which cells respond to their mechanical cues is mediated by FAK for both cell velocity and persistence.

FAK’s role in translating mechanical cues into biochemical migration responses is well established.^9,20,25,58^ FAK-null fibroblasts prefer to migrate away from softer substrates (due to generating less traction forces), and also demonstrate less directional persistence than controls.^20^ However, in a homogenous system with unchanging substrate stiffness, FAK inhibition has been shown to decrease migration dynamics by altering β1 mediated activation of FAK.^9^ Too much persistence has been associated with exacerbation of the fibrotic response due to promotion from pro-fibrotic signaling.^9^ Therefore, reduced migratory persistence is a mechanism by which FAK inhibition can decrease the fibrotic response.^9^ Our results suggest that when FAK is inhibited, cells no longer alter their persistent migration behavior in response to mechanical stimuli (substrate stiffnesses). Yet, FAKI-treated cells closed wounds faster on softer substrates mimicking normal wounds than FAKI-treated cells on stiffer substrates mimicking fibrotic wounds. Cells do not migrate in a directionless manner, therefore, we next studied the angle of cell migration during wound closure, a parameter assigning a vector to our directionality data to understand how the cells are angled toward the axis of wound closure during migration.

### FAK inhibition dysregulates angle of wound closure

The angle at which the end point of the cell’s trajectory points away from the origin (Fig S2B) indicates the orientation of the cell’s movement. The 0-180° x-axis, which is perpendicular to the wound along the x-axis, is logically the angle of most efficient wound closure. Based on our data (Fig 4A,C), fibroblasts in the DMSO control condition angle toward the x-axis, perpendicular to the wound edges, indicating that 180° is the angle oriented towards wound closure regardless of substrate stiffness. FAK inhibition dysregulates cell angle significantly away from the 0-180° wound closure axis regardless of substrate stiffness.

Therefore, cells are unable to orient themselves properly towards the wound when FAK is inhibited. Our data indicates that cells naturally close the wound at a rate independent of substrate stiffness, and suggests that when FAK is inhibited, the cell velocity, persistence, and angle are all significantly dysregulated, significantly slowing wound closure.

## Conclusions and Future Works

Our study shows that FAK mediates substrate stiffness induced fibroblast migration dynamics underlying wound closure. FAK is active at a particularly critical junction in the conversion of physical stimuli into biochemical cues – therefore, it appears that FAK is the critical element in regulating the cells’ ability to decipher and respond to substrate stiffness. As our data shows, without FAK, cells are unable to migrate rapidly.^59^ Previously, inhibition of FAK was demonstrated to increase actin contractile viscosity.^60^ FAK inhibition was also demonstrated to alter integrin expression, which in turn limited cell migration in response to extracellular matrix cues.^61^ FAK is an essential part of the focal adhesion complex maturation.^62^ Focal adhesions are stronger on stiffer substrates.^63^ Stronger focal adhesions allow for more nuclear elongation.^64^ Nuclear elongation contributes to efficacy of Yes associated protein (YAP) signaling by opening nuclear pores, which allows for more YAP nuclear translocation, and promotes migration.^65^ Additionally, greater recognition of the importance of group migration dynamics is increasingly recognized to impact wound closure dynamics. One emergent property is actin supra-structures^45^ which arises from the mechanotransduction effect on migration. Visual examination of fibroblast behavior in the FAK inhibited conditions (Fig 2A) indicates that the number of breakage events, or cells switching from group migration to individual migration, is greater than in the control conditions. This suggests that FAK is responsible for more than local substrate stiffness sensing, but also neighbor cell sensing mechanisms as well. FAK-YAP signaling is already known to be sensitive to the neighbor fraction of local cells, likely through integrin-associated activation, which can recognize both ECM and cells.^19^ Therefore, group migration, including multiple phenomenon such as breakage events or single cells breaking away from the migrating sheet and cell cluster tensions, could feed emergent properties that also affect wound closure. Moreover, FAK, YAP are known to have a complex relationship, leading to both regeneration and fibrosis. The field faces the question of how to tease apart this complicated relationship between FAK and YAP in the context of wound closure. Future directions might be well served in contextualizing FAK analysis with localized cell numbers and cell-cell connections.

Therefore, future directions should include the study of traction force microscopy to investigate if the decreased and impaired migratory phenotype during wound closure caused by FAK inhibition causes a mechanobiological switch towards a less contractile phenotype. Wound contraction plays an essential role in wound closure and surprisingly, it also decides wound fate, either regeneration or repair. Our results, contextualized within the literature, highlight FAK as a key regulator of cell migration in response to substrate stiffness. Fibroblasts on soft substrates mimicking normal wounds migrate slower and more randomly than fibroblasts on stiff substrates mimicking fibrotic wounds. However, when FAK is inhibited, fibroblasts cannot respond to differences in substrate stiffnesses. Instead, FAK inhibited fibroblasts migrate much slower, more randomly, and less angled toward wound closure, resulting in impaired wound closure rates. Here we show FAK is responsible for wound closure by controlling cellular migration dynamics (velocity, persistence, and angle of closure) in response to substrate stiffnesses mimicking normal or fibrotic wounds.

## Acknowledgments

K.W. acknowledges startup funds from the Bioengineering Department at Temple University, an Office of the Vice President for Research Bridge grant, and a grant (#2305) from the W.W. Smith Charitable Trust. N.A was supported by the Fulbright Foundation. J.D.B and A.O. were supported by the National Institute of Health (5T34 GM 136494). The authors thank the shared departmental resources at Temple University for equipment usage. We thank BioRender for providing a platform to create the schematics used in figures.

## Credit Author Statement

Jennifer Patten: Conceptualization, Methodology, Validation, Data Analysis, Visualization, Writing. Nourhan Albeltagy: Methodology, Validation, Data Analysis, Review and Editing. Jacob Bonadio: Methodology, Validation. Armando Ortez: Review and Editing. Karin Wang: Conceptualization, Methodology, Validation, Visualization, Writing, Funding acquisition, Supervision.

