## Supplementary Figures for "FAK Differentially Mechanoregulates Cell Migration During Wound Closure"

A

| Condition | Stiffness | Wound Closure (Best slope fit ± Slope Std Error) | Velocity (µm/min) | Directionality (0,1) | Distance (µm) | Angle (degrees) | Nuclear Aspect Ratio 48h (µm) |
|---|---|---|---|---|---|---|---|
| Media | 18 kPa | -2.022±0.277 | 0.26±0.1107 | 0.5563±0.1945 | 160.0±138.1 | 180.9±100.8 | 1.341±0.1484 |
| | 146 kPa | -1.939±0.3105 | 0.3328±0.1524 | 0.5983±0.1921 | 241.6±216.2 | 180.9±113 | 1.368±0.2333 |

B

| Condition | Stiffness | Wound Closure (Best slope fit ± Slope Std Error) | Velocity (µm/min) | Directionality (0,1) | Distance (µm) | Angle (degrees) | Nuclear Aspect Ratio 48h (µm) |
|---|---|---|---|---|---|---|---|
| DMSO | 18 kPa | -1.960±0.2762 | 0.2622±0.1004 | 0.5915±0.1731 | 168.0±150.6 | 176.4±111.5 | 0.1301±0.1402 |
| | 146 kPa | -1.979±0.3273 | 0.2977±0.1654 | 0.5117±0.2640 | 157.7±154.5 | 174.4±106.9 | 1.371±0.1778 |
| FAKI | 18 kPa | -1.924±0.2740 | 0.1531±0.109 | 0.4139±0.2049 | 125.7±108.6 | 164.5±104.8 | 1.311±1.525 |
| | 146 kPa | -1.396±0.2740 | 0.1752±0.2399 | 0.4278±0.2232 | 117.8±92.58 | 160.5±113.4 | 1.3731.311±1.525 0.1597 |

**Figure S1. Mean ± standard deviation for all data analyzed. A.** Data for media only condition corresponding to Figure 1. **B.** Comparison of all conditions corresponding to Figures 2-4 and Figure S2.

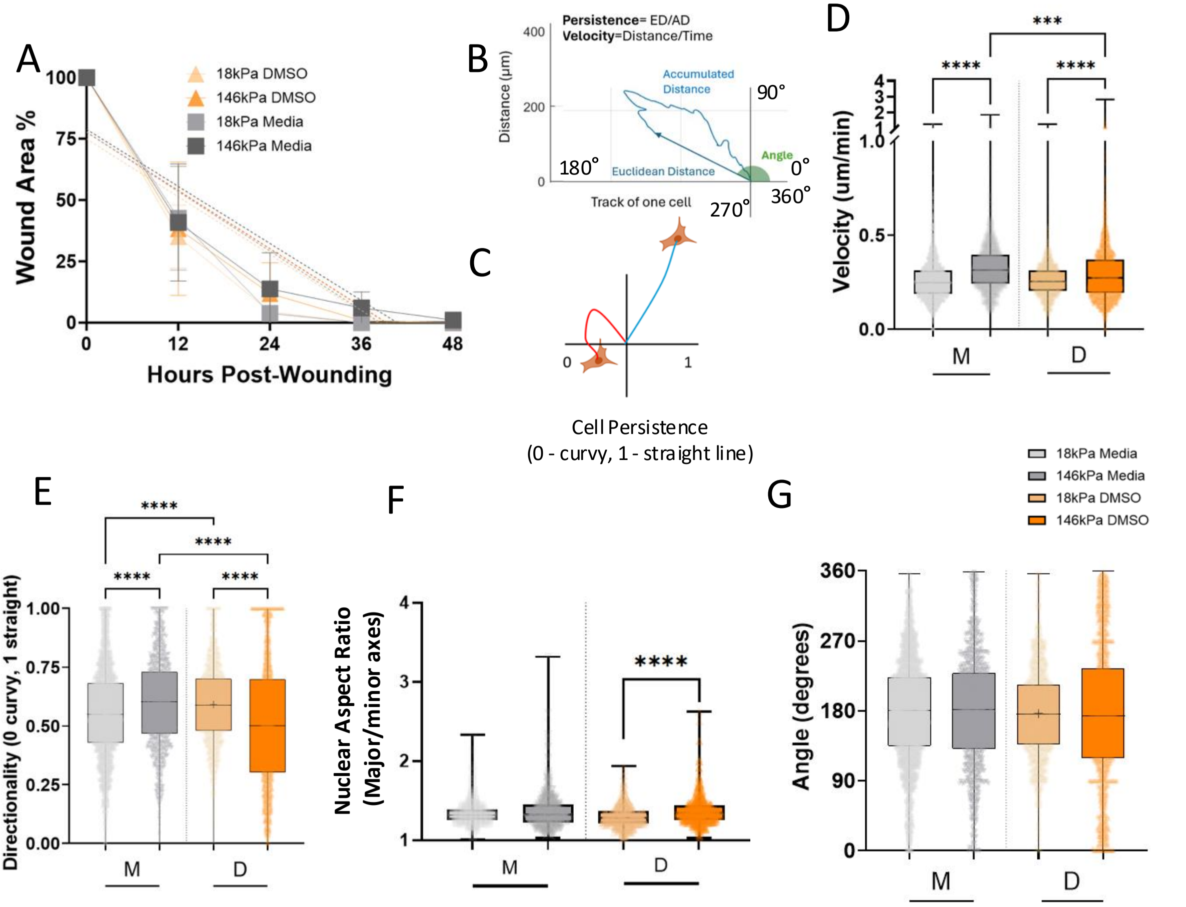


**Figure S2. Media comparison to DMSO controls. A.** Wound closure comparison between media and DMSO controls showed no significant differences. **B**. Schematic of persistence, where Persistence = Euclidean distance over accumulated distance. **C.** Schematic of persistence quantification, with 0 indicating random migration and 1 a continuous line. **D**. Comparison of media only conditions to DMSO controls indicated DMSO significantly slowed cell velocity on stiffer substrates, ***P=0.0006, ****P<0.0001. **E**. Comparison of media only condition to DMSO control for cell directionality indicates DMSO significantly upregulates directionality on soft substrates while downregulating directionality on stiffer substrates, ****P<0.0001. **F**. Comparison of media and DMSO controls indicate on the softer substrate stiffness, DMSO treatment decreases aspect ratio, turning the nuclei into a more circular shape corresponding to a less migratory shape, ****P<0.0001. **G**. Migration angle comparing DMSO to media. All N≥3 replicates. Two-way ANOVA, with Šídák's multiple comparisons post-hoc testing.

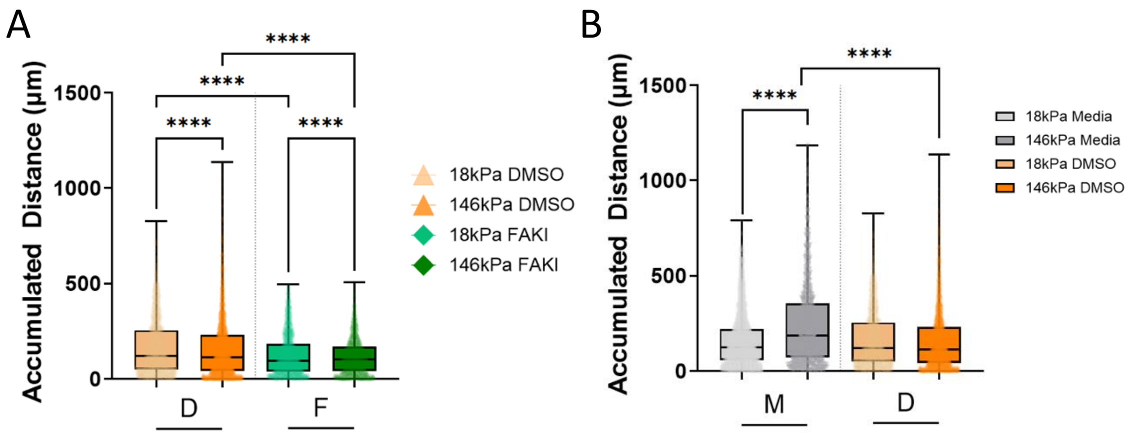


**Figure S3. Accumulated cell distance with comparison of media-only and DMSO controls. A.** Accumulated cell distance, ****P<0.0001. **B.** Comparison of media-only conditions with DMSO controls. ****P<0.0001. All N≥3 replicates. Two-way ANOVA, with Šídák's multiple comparisons post-hoc testing.

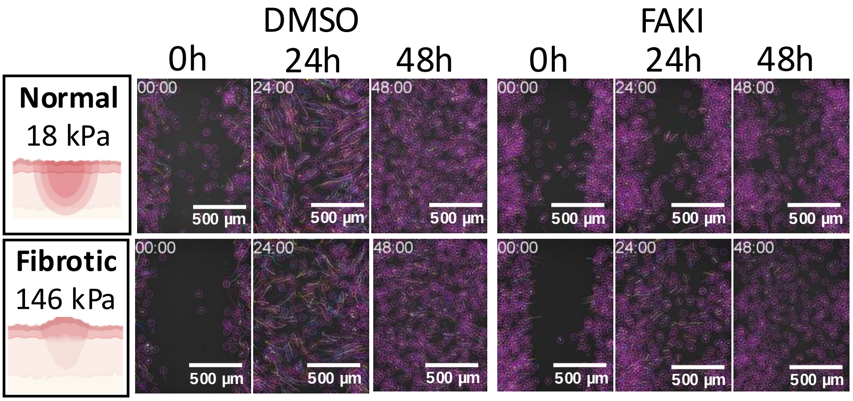


**Figure S4 Cell Dynamics** Representative Trackmate images at 0, 24, 48h post-wounding**.** Cell spots marked in purple circles, with cell tracks marked in colored lines**.** Scale bar 500um.

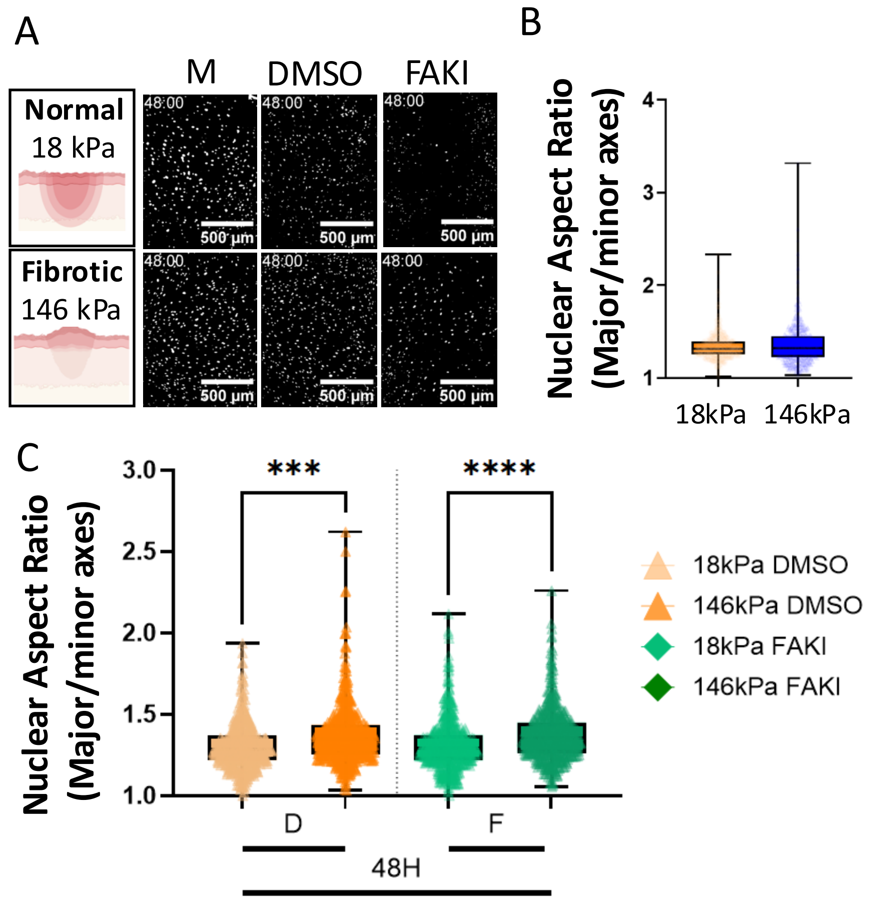


**Figure S5. FAK is not essential in regulating nuclear shape dynamics. A**. Representative 10x objective fluorescent images of cell nuclei (labeled with Spy595) were run through CellProfiler nuclei segmentation analysis to identify nuclei aspect ratio. **B**. Wound substrate stiffness does not alter nuclear shape during wound closure when inhibitors are absent in the media only condition. **C**. Mechanical substrate stiffness, regardless of FAK inhibition, regulates nuclear aspect ratio; cells on stiffer substrates mimicking fibrotic conditions are more elongated, indicating a more migratory phenotype than the cells on softer substrates mimicking normal wound stiffnesses, ***P=0.0002, ****P<0.0001. Two-way ANOVA, Šídák's multiple comparisons post-hoc analysis. N≥3 replicates.
